# Therapeutic Targeting of TIM-4-L With Engineered T Cells for Acute Myeloid Leukemia

**DOI:** 10.1101/2023.10.03.560752

**Authors:** Brandon Cieniewicz, Edson Oliveira, Mike Saxton, Damoun Torabi, Ankit Bhatta, Phanidhar Kukutla, Alexander Arballo, Zhou Yang, Bi Yu, Maria Fate, Hongxiu Ning, Lawrence Corey, Abhishek Maiti, Daniel Corey

**Affiliations:** Cero Therapeutics Inc., South San Francisco, CA 94080; Vaccine and Infectious Disease Division, 2 Fred Hutchinson Cancer Research Center, Seattle, WA 98109, USA; Department of Leukemia, University of Texas MD Anderson Cancer Center, Houston, TX 77030, USA

## Abstract

Disruption of the lipid asymmetric bilayer is a common feature observed in cancer cells. We utilized the natural immune receptor TIM-4 to interrogate for loss of plasma membrane phospholipid polarity in primary acute myelogenous leukemia (AML) samples. We performed FACs analysis in 33 patients and correlated TIM-4-L expression frequency and intensity with molecular disease characteristics. In normal tissues, TIM-4-L is confined to the internal leaflet of the plasma membrane. By contrast, 86% of untreated AML blasts in our analysis displayed upregulation of cell surface TIM-4-L. These observations were agnostic to AML genetic classification, as samples with mutations in *TP53, ASXL1*, and *RUNX1*, also displayed TIM-4-L upregulation similar to that seen in favorable and intermediate subtypes. This TIM-4-L dysregulation was also stably present in both Kasumi-1 and MV-4-11 AML cell lines. To evaluate the potential of upregulated TIM-4-L to serve as a target for adoptive T cell therapy (ACT), we constructed TIM-4-L-directed engineered T cells, which demonstrated potent anti-leukemic effects, effectively eliminating AML cell lines both in vitro and in vivo. This approach led to the eradication of AML cells across a range of endogenous TIM-4-L expression levels. These results highlight TIM-4-L as a highly prevalent and novel target for T cell-based therapy in AML. Further investigations into the role of TIM-4-L in AML pathogenesis and its potential as an anti-leukemic target for clinical development are warranted.

## Introduction

Development of engineered T cell therapies for acute myeloid leukemia (AML), a clinically and biologically heterogenous disease, has proven difficult in part due to the identification of suitable target antigens (1-4). Epigenetic profiling has revealed diverse alterations in AML (5), notably the dysregulation of key lipid transport molecules that drive immunosuppression and maintain lipid composition and structure within plasma membranes (6). Asymmetric lipid distribution in the plasma membrane is one of the fundamental features of living cells and changes to membrane organization have been observed in both virally-infected cells and hematologic and solid tumors (7-12). Normally sequestered to the inner leaflet, the externalization of phosphatidylserine promotes phagocytic clearance, facilitating the removal of cells to maintain tissue homeostasis (13). In cancer tissues, the effects of abnormal phospholipid distribution have been linked to varying, context-dependent functions. In advanced tumors, phospho-lipid signaling contributes to pathogenesis by shaping the tumor-promoting function of monocyte-derived tumor associated macrophages (TAM) (14, 15), whereas in nascent tumors, specific subsets of tissue-resident dendritic cells expressing high levels of the phosphatidylserine receptor TIM-4 lead to enhanced tumor antigen cross-presentation and the induction of cytotoxic T cell (CTL) anti-tumor responses (16).

These data led us to develop TIM-4 chimeric T cell-based immunotherapy against its native ligand phosphatidylserine, hereafter referred to as TIM-4-L. This therapeutic approach specifically targets cancer cells for cytotoxic and antigen presentation-mediated elimination driven by dysregulation of cell surface TIM-4-L (17). We previously demonstrated upregulation of TIM-4-L in B cell malignancy primary specimens and cell lines and developed a therapeutic approach to achieve lymphoma control by eliciting both direct cytotoxic effects and indirect-mediated cross priming (17). In this report, we use the natural immune receptor TIM-4 to interrogate lipid distribution in primary AML specimens and cell lines and characterize the anti-leukemic effects of TIM-4-L-targeting T cells in AML cell line models. Previous analyses of acute promyelocytic leukemia cells demonstrated that lipid scrambling contributes to disease pathogenesis by forming a scaffold for blood-clotting factors, ultimately leading to thrombosis (9). Similarly, ATP11A downregulation, responsible for membrane phospholipid internalization, correlates with worse outcomes in AML subtypes (6).

To elucidate the pattern of TIM-4-L on AML we performed a detailed FACs analysis in 33 patients and correlated expression frequency and intensity with molecular disease characteristics. In normal tissues, TIM-4-L is confined to the internal leaflet of the plasma membrane. By contrast, 86% of untreated AMLs in our analysis displayed upregulation of cell surface TIM-4-L. Importantly, AMLs with adverse genetic features, such as mutations in *TP53, ASXL1*, or *RUNX1*, also displayed TIM-4-L upregulation similar to that seen in favorable and intermediate subtypes. These findings were replicated in established AML cell lines with diverse genetic alterations, all of which exhibited upregulation of TIM-4-L. We further establish that TIM-4-L-targeting T cells are highly effective in eliminating AML cell lines in xenograft mouse models. Taken together, the results obtained in these studies demonstrate that TIM-4-L is a viable target for T cell therapy in AML and support additional preclinical and clinical development.

## Materials and Methods

### Patient Samples

Patient-derived AML bone marrow and PBMC samples and healthy donor controls were obtained from a contract research organization (Discovery Life Sciences). Samples were collected under IRB and ethics committee approval. All patients gave written informed consent to participate. Donor information is listed in **Table 1** and **Supplementary Table 1**.

**Table 1.** Patient Characteristics.

| Characteristic | Number (%) of Patients (n=33) |
| --- | --- |
| **Age, years** |  |
| Mean | 58.6 |
| Range | 21-88 |
| **Sex** |  |
| Male | 15 (45%) |
| Female | 18 (55%) |
| **Newly Diagnosed** | 87.9% |
| **AML, marrow blasts, mean** | 75.8% |
| Range | 15-95% |
| **Risk Stratification** |  |
| Adverse | 21 (63.6%) |
| Intermediate | 6 (18.2%) |
| Favorable | 1 (3.0%) |
| Unknown | 5 (15.2%) |
| **Genetic abnormality<sup>a</sup>** |  |
| ASXL1 | 30.3% |
| BCOR | 30.3% |
| EZH2 | 9.1% |
| RUNX1 | 21.2% |
| SF3B1 | 6.1% |
| SRSF2 | 12.1% |
| STAG2 | 15.2% |
| U2AF1 | 0.0% |
| ZRSR2 | 15.2% |

<sup>a</sup> adverse risk mutations, ELN 2022 risk classification by genetics at initial diagnosis

### AML Cell Lines and Reagents

Kasumi-1 (ATCC, Cat# CRL-2724), MV-4-11 (ATCC, Cat# CRL-9591), and THP-1 cells (ATCC, Cat# TIB-202) were cultured in recommended media. To generate stable mCherry+ cells for use in co-culture assays, Kasumi-1 or MV-4-11 cells were transduced with Incucyte Nuclight Red lentivirus (Essen Bioscience, MI). Transduced cells were selected with puromycin and tested for purity by flow cytometry. Kasumi-1 cells that constitutively express firefly luciferase and GFP were purchased from Cellomics (Cat# SC-1152). 5-Azacytidine was obtained from MedKoo (Cat# 100070) and diluted into a stock solution in diH2O.

### Generation of Viral Vectors

Replication-defective lentiviruses were generated using standard methods, employing a third-generation lentiviral plasmid that encodes the extracellular domain of TIM-4, CD28 transmembrane domain, and CD28, CD3ζ, and TLR2/TIR-1 intracellular signaling domains (18). In brief, the lentiviral plasmid was combined with three packaging plasmids encoding VSV-G (Aldevron, Cat# 5037-5), gag/pol (Aldevron, Cat# 5035-5), and rev (Aldevron, Cat# 5033-5). This mixture was then transfected into HEK293T cells (ATCC, Cat# CRL-3216) cultured in LV-MAX™ Production Medium (ThermoFisher, Cat# A3583401) using polyethyleneimine linear (Polysciences, Cat# 23966-1). After 48 hours, the lentiviral vectors were harvested by centrifuging the cell culture at 400 x g at 4 °C to separate the supernatants containing the vectors. The supernatant was divided into aliquots ranging from 500 μL to 1 mL and stored at -80 °C until further use.

### Generation of TIM-4-chimeric T cells

GMP clinical manufacturing procedures were utilized in experiments. T cells were positively selected using anti-CD4 and CD8 antibodies through the Prodigy TCT program on a CliniMACS Prodigy (Miltenyi Biotec, Cat# 200-075-301). After selection, a fixed number of cells were used to start the TCT program on Prodigy using a TS520 tubing set (Miltenyi Biotec, Cat# 170-076-600). Cells were cultured in Optimizer medium (ThermoFisher, Cat# A1048503) and activated by using GMP grade MACS T cell TransAct (Miltenyi Biotec, Cat# 200-076-202). After activation, cells were transduced with lentiviral vector encoding the extracellular domain of TIM-4, CD28 transmembrane domain, and CD28, CD3ζ, and TLR2/TIR-1 intracellular signaling domains (CER-1236). After transduction, cells were washed and resuspended in fresh medium for expansion and medium was supplied or refreshed. At harvest day, cells were harvested into freshly prepared optimizer medium. After cell counting, cells were cryopreserved. T cell viability, expansion, transduction efficiency, VCN, PD-1 expression, T cell memory profile, CD4/8 ratio, and target-dependent IFN-γ secretion were measured at the end of production as previously described (Supplementary Figure 1) (17). All in vitro and in vivo animal studies were done with thawed cryopreserved TIM-4 chimeric T cells to mimic actual clinical conditions.

### Flow cytometry of AML samples

To assess cell surface TIM-4-L on primary samples, cryopreserved specimens were thawed at 37°C, then transferred to 9ml of cold media. All subsequent staining were performed with cold buffer and on ice. Cells were resuspended in TIM-4 binding buffer (PBS (ThermoFisher, Cat# 14190-144) + 0.5mM CaCl_2_ (VWR, Cat# E506)) and counted using a ViCell Blu automated cell counter (Beckman Coulter, Cat# C19196)). Cells were plated and stained using Live/Dead Fixable Aqua (Thermofisher, Cat# L34957)), CD45 (Clone HI30, Biolegend, Cat# 304052)), and recombinant human TIM-4-HIS (R&D Systems, Cat# 9407-TM-050)). After primary staining, cells were washed with TIM-4 binding buffer, then stained with anti-HIS tag secondary (Cell Signaling Technology, Cat# 14931S). Cells were washed with TIM-4 binding buffer, then samples were analyzed on a Cytoflex LX (Beckman Coulter, Cat# C40323). TIM-4-L expression was assessed in live, CD45^+^ cells using FlowJo v10 (BD Biosciences, v10). Gates for TIM-4-L positivity were set using a secondary only control sample.

### Co-culture cytotoxicity, cytokine secretion, and proliferation assays

CER-1236 T cells were co-cultured with engineered Kasumi-1 or MV-4-11 mCherry+ cells at 1:1 or 1:2 effector:target (E:T) ratios in cytokine-free media. Cell numbers were calculated using TIM-4^+^ cells for each T cell product. Cytotoxicity was monitored using the IncuCyte S3 Live-Cell Analysis System with images acquired every 2 h. Reduction of mCherry^+^ signal was used to determine target cell elimination. Untransduced T cells were used as negative controls. Parallel plates were prepared for supernatant collection at 120 h. IFN-γ secretion was measured using the ELLA automated immunoassay platform (Biotechne, Cat# ST01B-PS-002574). To assess T cell proliferation, co-culture samples were collected at 120h and stained for live/dead fixable Aqua and CD3 (Clone HIT3a, Biolegend, Cat# 300306) in FACS buffer. To enable cell enumeration, Precision Count Beads (Biolegend, Cat# 424902) were added to each sample. Total live CD3^+^ cells were estimated per sample and divided by the number of initially plated cells to determine fold expansion.

### Plate-bound TIM-4-L T cell stimulation assay

To generate plates coated with TIM-4-L, porcine brain-derived PS (Avanti Polar Lipids, Cat# 840032C) was used to coat plates with 0.5 or 5ug/ml TIM-4-L. The 0 μg/mL wells are treated with methanol and allowed to evaporate to serve as the uncoated unstimulated control. Each stock (20 μL) was added to a flat-bottomed 96-well plate and rocked to ensure uniform coverage of the surface, then allowed to evaporate with the lid off in a biosafety cabinet.

Cryopreserved CER-1236 T cells, or untransduced (UNT, control) T cells prepared were thawed and plated at 25,000 CER^+^ T cells per well, or equivalent numbers of UNT T cells. 5-Azacytidine was added to each well at 1, 10, 100, 1000, and 10000 nM. After incubation for 24 h, supernatants were collected and secretion of IFN-γ was measured using the ProteinSimple Ella automated immunoassay system. Duplicates were measured independently and averaged.

### In vivo assays

All experimental procedures and protocols were approved by the Institutional Animal Care and Use Committee of Charles River Labs (Wilmington, MA). Animals were kept in a pathogen-free environment with a filtered air supply. Five- to 7-week-old female NSG MHC dKO mice (NOD.Cg-Prkdcscid H2-K1b-tm1Bpe H2-Ab1g7-em1Mvw H2-D1b-tm1Bpe Il2rgtm1Wjl/SzJ) were obtained from Jackson Laboratory (Cat# 205216, RRID: IMSR_JAX:025216).

For a disseminated in vivo model of AML, NSG mice were engrafted i.v. via the lateral tail vein with 2e^6^ Kasumi-1 fLuc/GFP cells in 100 μL PBS without magnesium or chloride ions on day −12. Mice were randomized on day -1, then day 0 animals were infused i.v. via the lateral tail vein with 7.5e^6^ CER-1236 T cells in 100 μL PBS without magnesium or chloride ions. Control mice were infused i.v. with untransduced T cells equivalent to the dose given to the 7.5e^6^ CER-1236 T cell groups. Animals were assessed daily for signs of GvHD, including hair loss, skin rash, skin thickening, and hunched posture.

Tumor burden was monitored twice per week using BLI measured on an Ami HTX imager (Spectral Instruments, Tucson, AZ) following an intraperitoneal injection of IVISbrite D-luciferin substrate solution (Cat# 122799, 100 μL, 30 mg/mL). Body weight was measured weekly. Peripheral blood was collected weekly by submandibular bleed into a K2-EDTA tube (BD, Cat# 365974) and stored at −80°C.

### Pharmacokinetics analysis

Genomic DNA was isolated from peripheral blood samples using the Maxwell RSC Blood DNA kit (Promega, Cat# AS1400 using the Maxwell RSC benchtop automated DNA/RNA extraction instrument (Promega). Total CER-1236 T cells present in the blood was determined using ddPCR as previously described (17). The following equation was used to determine copy number per μg of DNA: (copies of CER-1236 per μL of reaction * 1000000)/(copies of HAT gene per μL of reaction * 3.3) (19).

### Statistical analysis

All experimental data are presented as mean ± standard error of the mean or standard deviation, as indicated in figure legends. Statistical significance between two groups was determined using two-tailed Student’s t test. For multiple group comparison, One- or two-way ANOVA was used. A p value of less than 0.05 was considered statistically significant. Data were analyzed and presented with GraphPad Prism.

## Results

### TIM-4-L Expression Across AML Molecular Subgroups

AML is characterized by its clinical and biological diversity. In a previous study, TIM-4-L expression was identified in a limited subset of AML cases, suggesting its involvement in disease pathogenesis (9). To delve deeper into these findings, we conducted an in-depth FACs analysis, quantifying surface TIM-4-L in 33 cryopreserved primary bone marrow aspirates from AML patients. Of these, 29 of 33 (87.8%) AML samples were taken from treatment-naïve patients, providing a baseline measurement of TIM-4-L prior to any treatment interventions. Our analysis revealed significantly higher cell surface TIM-4-L positivity and MFI compared to healthy donor counterparts (**Figure 1A, B**). AML blasts showed a range of TIM-4-L positivity, while healthy donor bone marrow aspirates showed only low levels of TIM-4-L positivity (**Figure 1A**). Notably, abundance of cell surface TIM-4-L, as assessed by gMFI, was significantly higher on AML blasts (**Figure 1B**) than on healthy donor bone marrow, which had a more uniform, overall low gMFI of TIM-4-L. TIM-4-L positivity was primarily observed in the CD45 intermediate AML population, as non-blast cells had low levels of TIM-4-L (**Supplementary Figure 2**).

**Figure 1.**
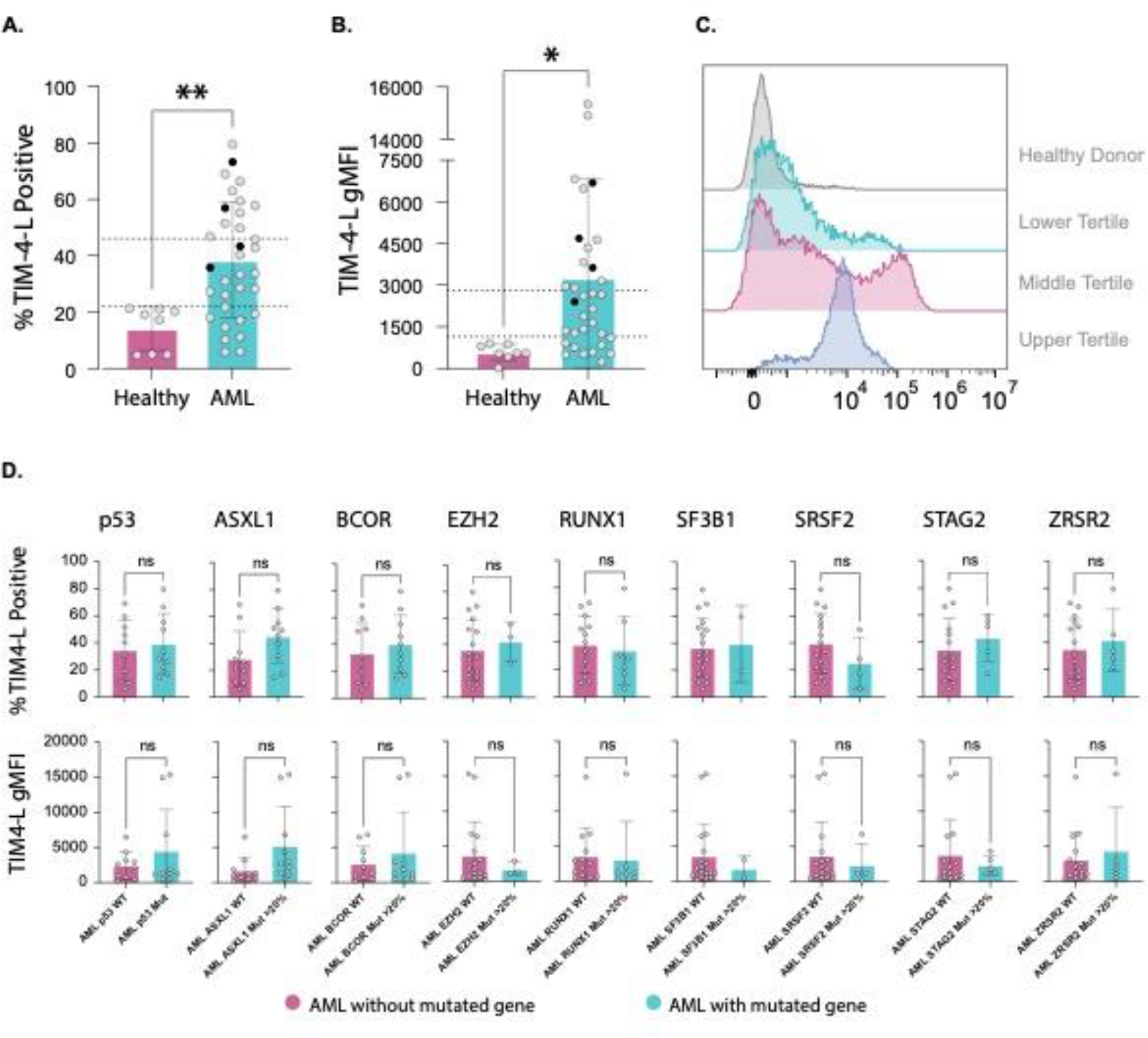
Quantification of TIM-4-L in Primary Tissues Across AML Molecular Subgroups. 33 primary AML bone marrow aspirates and 8 healthy donor bone marrow aspirates were stained for TIM-4-L using recombinant human TIM-4. (A) Percent positive TIM-4-L cells. Percent TIM-4-L-positive cells were identified on live, CD45dim cells. Gating was determined using TIM-4 FMO controls for each sample. Dotted lines indicate bottom, middle, and upper tertiles of TIM-4-L positivity. Black dots indicate samples receiving therapy. Each sample was stained in duplicate and average is shown. (B) gMFI of TIM-4-L staining. The gMFI of TIM-4-L staining was measured on live, CD45dim cells. Gating was determined using TIM-4 FMO controls for each sample. Dotted lines indicate bottom, middle, and upper tertiles of TIM-4-L positivity. Black dots indicate samples receiving therapy. (C) Representative flow images of AML from each tertile and healthy donor. (D) TIM-4-L positivity and staining intensity by adverse risk mutations. Newly diagnosed AML were grouped by VAF of >20 of the adverse risk mutations *ASXL1, BCOR, EZH2, RUNX1, SF3B1, SRSF2, STAG2*, or *ZRSR2* or >10 of *TP53*. (E) TIM-4-L positivity by AML risk group. Newly diagnosed AML bone marrow were grouped by risk group based on ELN 2022 risk classification parameters. Statistical significance was tested using student’s unpaired t-test. * = p < 0.05, ** = p < 0.01, ns = not significant.

Given the range of TIM-4-L observed on AML blasts, we separated them into 3 tertiles, corresponding to low, medium, and high TIM-4-L expression (**Figure 1A, B; dotted lines**). TIM-4-L positivity in the lowest tertile averaged 13.83% (n=9, range 5.88-21.8), which was similar to the 14.18% (n=8, range 4.94-19.1) TIM-4-L positivity observed in healthy bone marrow aspirates. The middle tertile had an average TIM-4-L positivity of 33.75% (n=10, range 26.1-45.9), representing a 2.44 fold increase over the lowest tertile. The highest tertile had an average TIM-4-L positivity of 59.92% (n=10, range 46.85-79.65), representing a 4.33 fold increase over the lowest tertile. Similar trends were observed in the TIM-4-L gMFI, with the lowest tertile having an average TIM-4-L gMFI of 665.3 (n=9, range 237.5-1103.5), comparable to the average of 582.3 (n=8, range 20.7-896) observed in healthy donor samples. The middle tertile of samples had an average TIM-4-L gMFI of 1897.1 (n=10, range 1257.5-2698), representing a 2.85 fold increase in per-cell TIM-4-L as compared to the first tertile. The upper tertile had an average TIM-4-L gMFI of 6538.4 (n=10, range 2908-15344), representing a 9.83 fold increase in per-cell TIM-4-L. Representative histograms for each tertile are shown in **Figure 1C**.

In addition to the treatment naïve patient samples, we measured TIM-4-L on 4 patients receiving various AML therapies (**Figure 1A, B, Black dots, Table 1**). These AML samples had overall high frequency of surface TIM-4-L. The per-cell TIM-4-L levels of the treated AML samples also grouped with the upper tertiles of treatment-naïve AML samples, as measured by gMFI.

Of the 33 bone marrow samples we tested, 18 were sequenced for variant alleles, allowing us to classify these samples according to adverse risk mutations based on the European LeukemiaNet (ELN) 2022 risk classification recommendations (20, 21). We compared TIM-4-L positivity and gMFI for *TP53, ASXL1, BCOR, EZH2, RUNX1, SF3B1, SRSF2, STAG2*, and *ZRSR2* mutations. There were no differences in TIM-4-L positivity or MFI on the basis of genetic status; TIM-4-L positivity and MFI were equally distributed among genetically high-risk patients (**Figure 1D**). When assessed more broadly by ELN risk categorization, we observed no differences in TIM-4-L positivity or gMFI between groups (**Supplementary Figure 3A, B**).

We also evaluated TIM-4-L the peripheral blast population in 10 primary samples and 4 healthy donor samples. Here we also noted higher TIM-4-L positivity in comparison to healthy donor PBMCs (**Supplementary Figure 4A**). Interestingly, although there was a trend towards increased per-cell TIM-4-L on circulating AML blasts, this difference did not reach statistical significance (**Supplementary Figure 4B**).

### Upregulation of Endogenous TIM-4-L in AML Cell Lines Kasumi-1 and MV-4-11 Cells and Evaluation of TIM-4-L Targeted Cell Therapy

Given the range of TIM-4-L positivity we observed in primary AML samples, we sought to determine if AML cell lines had similar endogenous levels of TIM-4-L expression. We assessed TIM-4-L on the AML cell lines Kasumi-1 and MV-4-11, harboring mutations in *TP53* and *FLT3* respectively. Interestingly, we noted relatively lower levels of surface TIM-4-L on these cell lines, with MV-4-11 TIM-4-L gMFI of 534 and Kasumi-1 a gMFI of 2363 (**Figure 2A**), corresponding to the lower and middle tertiles of TIM-4-L observed in primary samples.

**Figure 2.**
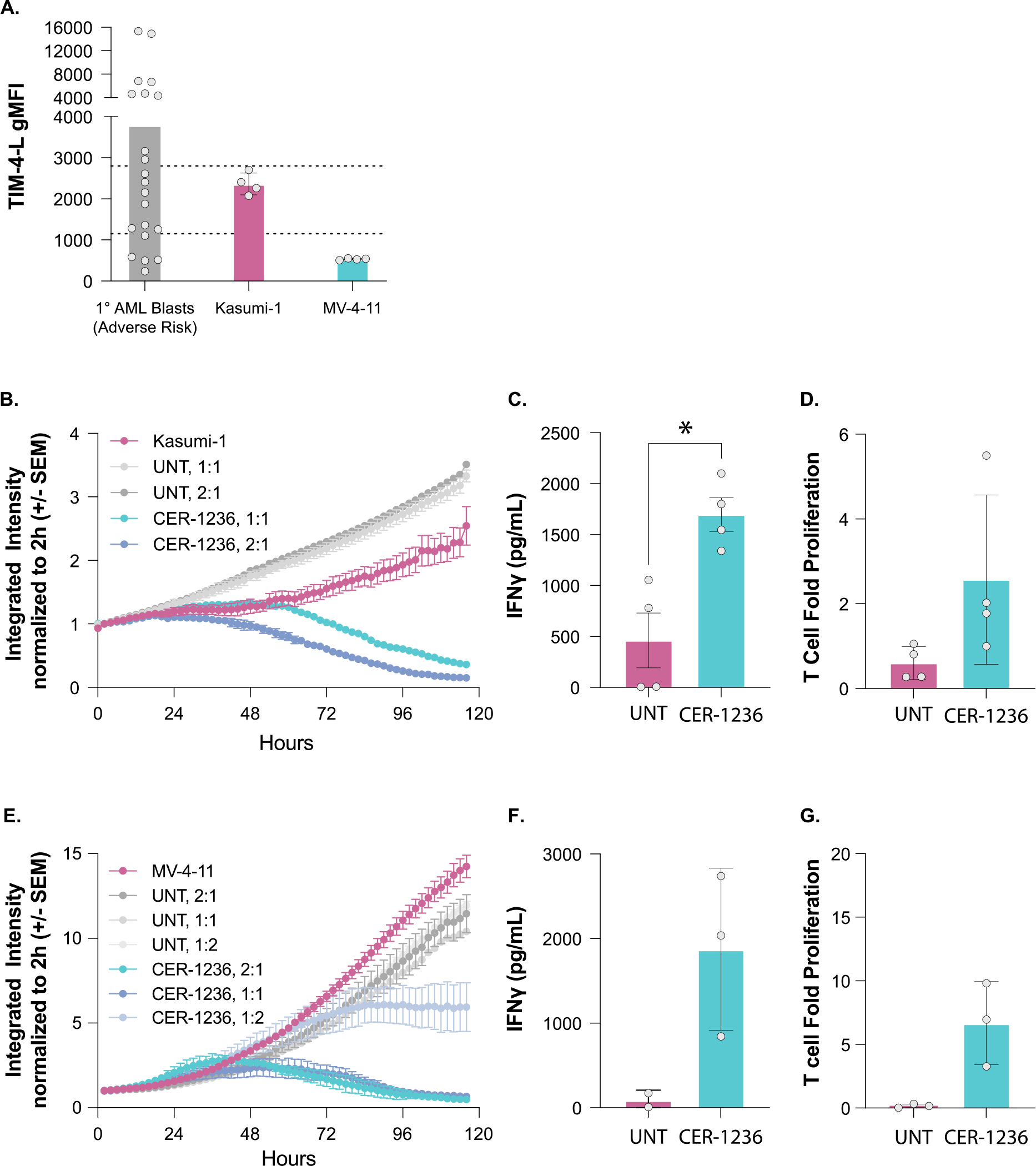
CER-1236 T cells eliminate the AML cell lines Kasumi-1 and MV-4-11 in vitro. (A) TIM-4-L on AML cell lines. The AML cell lines Kasumi-1 and MV-4-11 were stained for TIM-4-L using recombinant human TIM-4. TIM-4-L gMFI was measured in live cells and compared to TIM-4-L of adverse risk primary AML blasts. Average ± SEM is shown for each cell line, N=4. (B) CER-1236-mediated cytotoxicity against Kasumi-1 AML. Kasumi-1 mCherry^+^ cells were co-cultured with CER-1236 cells at a 2:1 or 1:1 E:T ratio and cytotoxicity was monitored for 120h. Average of 2 duplicate measurements is shown. Data is representative of n=4 experiments. (C) IFN-γ secretion in co-culture. Supernatant was collected from 2:1 E:T co-cultures at 120h and IFN-γ secretion was measured. Data points represent average of 2 duplicate measurements, n=4. (D) T cell proliferation in co-culture. After 120h of co-culture at 2:1 E:T ratio, total T cells in culture was determined using flow cytometry. Fold proliferation was calculated by dividing calculated cells at the end of co-culture by the number of total T cells plated at 0h. Data points represent average of 2 duplicate measurements, n=4. (E) CER-1236-mediated cytotoxicity against MV-4-11 AML. MV-4-11 mCherry^+^ cells were co-cultured with CER-1236 cells at a 2:1, 1:1, or 0.5:1 E:T ratio and cytotoxicity was monitored for 120h. Average of 2 duplicate measurements is shown. Data is representative of n=3 experiments. (C) IFN-γ secretion in co-culture. Supernatant was collected from 2:1 E:T co-cultures at 120h and IFN-γ secretion was measured. Data points represent average of 2 duplicate measurements, n=3. (D) T cell proliferation in co-culture. After 120h of co-culture at 2:1 E:T ratio, total T cells in culture was determined using flow cytometry. Fold proliferation was calculated by dividing calculated cells at the end of co-culture by the number of total T cells plated at 0h. Data points represent average of 2 duplicate measurements, n=3.

We next evaluated the therapeutic potential of targeting TIM-4-L by generating TIM-4-L directed engineered T cells known as CER-1236. CER-1236 T cells are engineered to express a chimeric receptor consisting of a TIM-4 extracellular domain fused to a CD28 transmembrane domain, and CD28, CD3ζ, and TLR2 intracellular signaling domains. This approach combines TIM-4 with T cell activation and innate signaling domains, leveraging the phagocytic and TIM-4-L-binding properties of TIM-4, the pro-phagocytic functions of TLR2, and the T cell activation capacity of CD3ζ and CD28. Cells were generated using lentiviral transduction of CD4^+^/CD8^+^ selected healthy donor T cells in the CliniMACs Prodigy. T cell expansion, transduction efficiency, VCN, memory subsets, and response to plate-bound TIM-4-L were comparable to previous reports (**Supplementary Figure 1)** (17). Previous pre-clinical studies have demonstrated the effective elimination of patient-derived lymphoma and non-small lung cancer xenografts by CER-12336 (17). In co-culture studies against MV-4-11 and Kasumi-1 cells, we measured target-dependent cytotoxicity, cytokine secretion, and T cell proliferation. CER-1236 T cells demonstrated strong cytotoxic responses against Kasumi-1 cells, which exhibited the higher endogenous TIM-4-L gMFI of the tested AML cell lines (**Figure 2B**). To further evaluate target specificity, we measured cytokine production and cell proliferation of CER-1236 and unmodified control T cells following coculture with target cells. Significant increases in IFN-γ secretion were measured in CER-1236 T cells compared to unmodified control T cells (**Figure 2C**). Additionally, modest levels of CER-1236 proliferation were measured in co-culture studies with Kasumi-1 cells (**Figure 2D**), although these did not reaching statistical significance.

We then tested CER-1236 responses against MV-4-11 AML cells, which harbor a *FLT3*-ITD mutation and exhibit lower overall TIM-4-L compared to Kasumi-1 cells. Even at low endogenous TIM-4-L expression levels, CER-1236 T cells displayed robust cytotoxicity against MV-4-11 cells, resulting in complete elimination of targets by 120h of co-culture (**Figure 2E**). This cytotoxic activity coincided with increased IFN-γ secretion (**Figure 2F**) and induction of T cell proliferation (**Figure 2G**).

Collectively, these data indicate highly specific reactivity by TIM-4-L-targeting T cells against AML cell lines expressing endogenous levels of TIM-4-L.

### Addition of the Hypomethylating Agent 5-azacytidine Does not Impair CER-1236 T cell Responses

5-azacytidine is a hypomethylating agent used commonly for newly diagnosed AML patients and as maintenance therapy for patients in remission (20, 21). Given previous observations that 5-azacytidine treatment of T cells can impair effector functions and mediate immunomodulatory effects (22-24), we sought to determine if concurrent treatment with 5-azacytidine affected CER-1236 T cell effector functions. We repeated co-culture experiments with Kasumi-1 and MV-4-11 AML cell lines in the presence or absence of 100nM of 5-azacytidine, which corresponds to nearly half of Cmax with standard IV/SQ azacytidine dosing (25)**(Figure 3A)**. After 5 days of co-culture, we assessed specific killing of either cell line. CER-1236 T cells eliminated 83.1% of Kasumi-1 cells over the course of the co-culture experiment (**Figure 3B**). The addition of 5-azacytidine had no significant effect on CER-1236 cytotoxicity, with 83.0% of Kasumi-1 eliminated in the presence of drug. The addition of 5-azacytidine had no deleterious effect on the elimination of MV-4-11 cells either, with 72.3% of target cells eliminated in the absence of drug as compared to 83.3% of cells eliminated in the presence of drug (**Figure 3C**). 100nM of 5-azacytidine alone had only minor effects on Kasumi-1 cell growth. However, we observed that treatment of both Kasumi-1 and MV-4-11 cells with 5-azacytidine could induce TIM-4-L **(Supplementary Figure 5)**.

**Figure 3.**
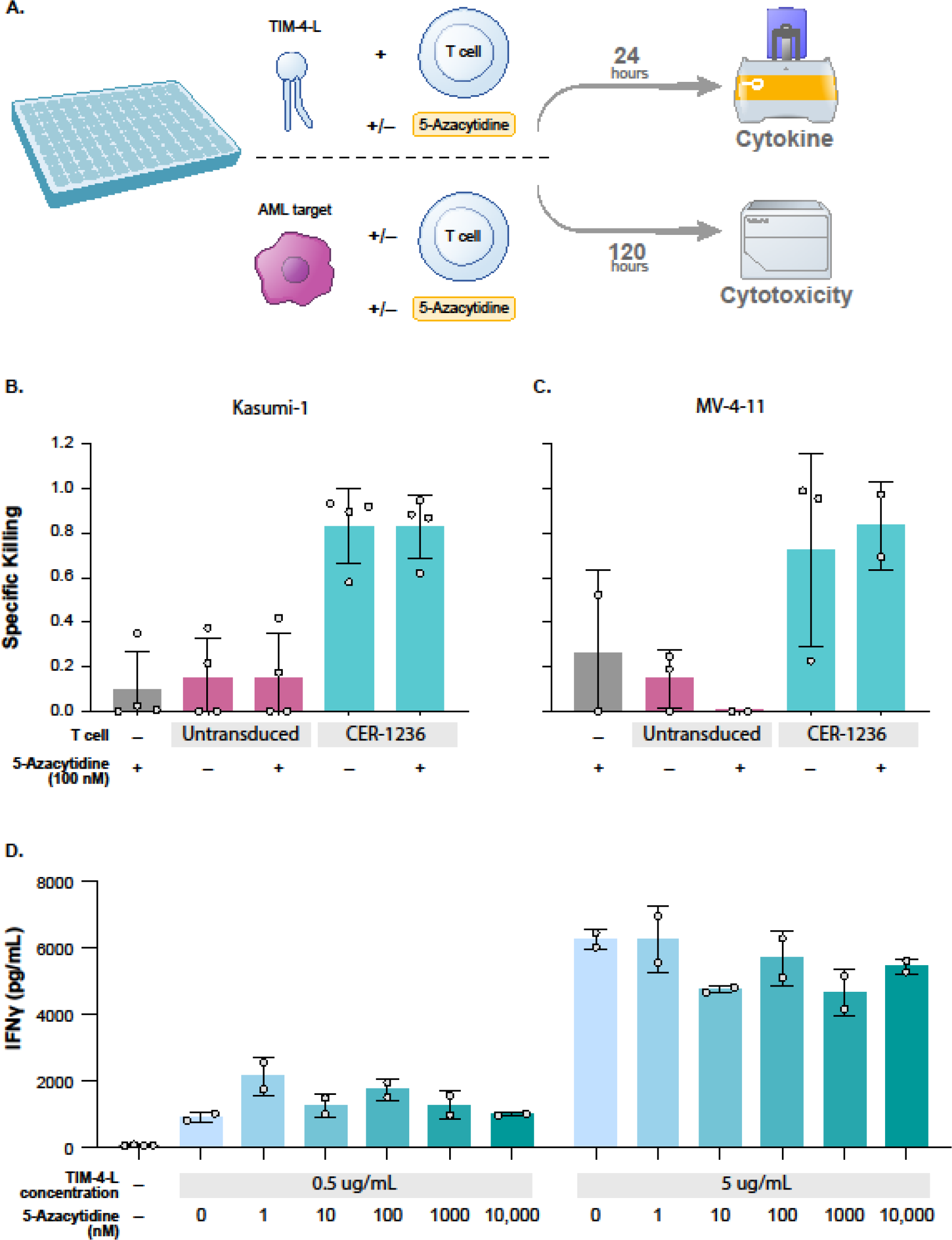
5-Azacytidine does not impair CER-1236 T cell function in co-culture. (A) Schematic of co-culture or plate-bound TIM-4-L stimulation assay with 5-azacytidine. (B,) 5-azacytidine effect on CER-1236-mediated cytotoxicity. CER-1236 and Kasumi-1 mCherry^+^ cells (B) or MV-4-11 (C) were co-cultured at a 2:1 E:T ratio in the presence or absence of 100nM 5-azacytidine. After 120h, the specific killing was determined for each condition using the equation 1-(integrated intensity for treated sample/integrated intensity for Kasumi-1 mCherry^+^ cells alone). Data points represent average of 2 duplicate measurements, n=4 for Kasumi-1 and n=3 for MV-4-11. (D) 5-azacytidine effect on TIM-4-L-mediated stimulation of CER-1236. CER-1236 T cells were stimulated on plates coated with 0.5 or 5μg/ml of TIM-4-L in the presence of 1-10000nM of 5-azacytidine. After 24h, supernatant was collected and secretion of IFN-γ was measured. Average of 2 duplicate measurements is shown. Data is representative of n=2 experiments. Statistical significance was determined using a one-way ANOVA with Tukey’s post-test. * = p < 0.05, ns = not significant.

To further characterize the effect of 5-azacytidine on CER-1236 T cell function, we stimulated CER-1236 T cells with plate-bound TIM-4-L in the presence of a range of 5-azacytidine concentrations and measured IFN-γ secretion at 24h. 5-Azacytidine concentrations of up to 10μM had no effect on CER-1236 T cell responses to high or low concentrations of TIM-4-L (**Figure 3D**), indicating that CER-126 T cells remain functional across a range of 5-azacytidine concentrations.

### TIM-4-L Targeting T Cells Demonstrate Potent Target-Specific Cytotoxicity Against Kasumi-1 cells In Vivo

Given the potent anti-tumor response we observed in vitro, we sought to determine if CER-1236 T cells could eliminate Kasumi-1 xenografts in vivo. While mice have not been feasible models for assessment of CAR T safety due to differences in target antigens between species, this is not the case for CER-1236 T cells. TIM-4-L is conserved between mice and humans, allowing for potential on-target off tumor evaluation of potential toxicities. Using Kasumi-1 Fluc/GFP NSG MHC dKO xenografts, our initial assessment of TIM-4-L on leukemic cells demonstrated notably elevated levels of expression (**Figure 4A**). In healthy mouse cells no detectable TIM-4-L expression was observed. Established xenografts were subsequently treated with 7.5e6 CER-1236 T cells or an equivalent number of untransduced control T cells (**Figure 4B**). BLI was monitored weekly to assess tumor expansion and anti-tumor effects, while body weight was measured to assess animal health. Treatment with CER-1236 T cells mediated a potent anti-tumor effect, leading to elimination of tumor in all treated animals (**Figure 4C, D**). In contrast, animals that received untransduced T cells all exhibited disease progression. Consistent with enhanced antitumor activity, CER-1236 T cells rapidly expanded in the peripheral blood, peaking 7 days after T cell administration (**Figure 4E**) and gradually contracted over the course of 4-week period. Importantly, animals treated with CER-1236 T cells exhibited no reduction in body weight (**Figure 4F**) or signs of morbidity over the course of the experiment, underscoring the specific targeting of AML tumor cells by CER-1236 T while sparing healthy mouse cells.

**Figure 4.**
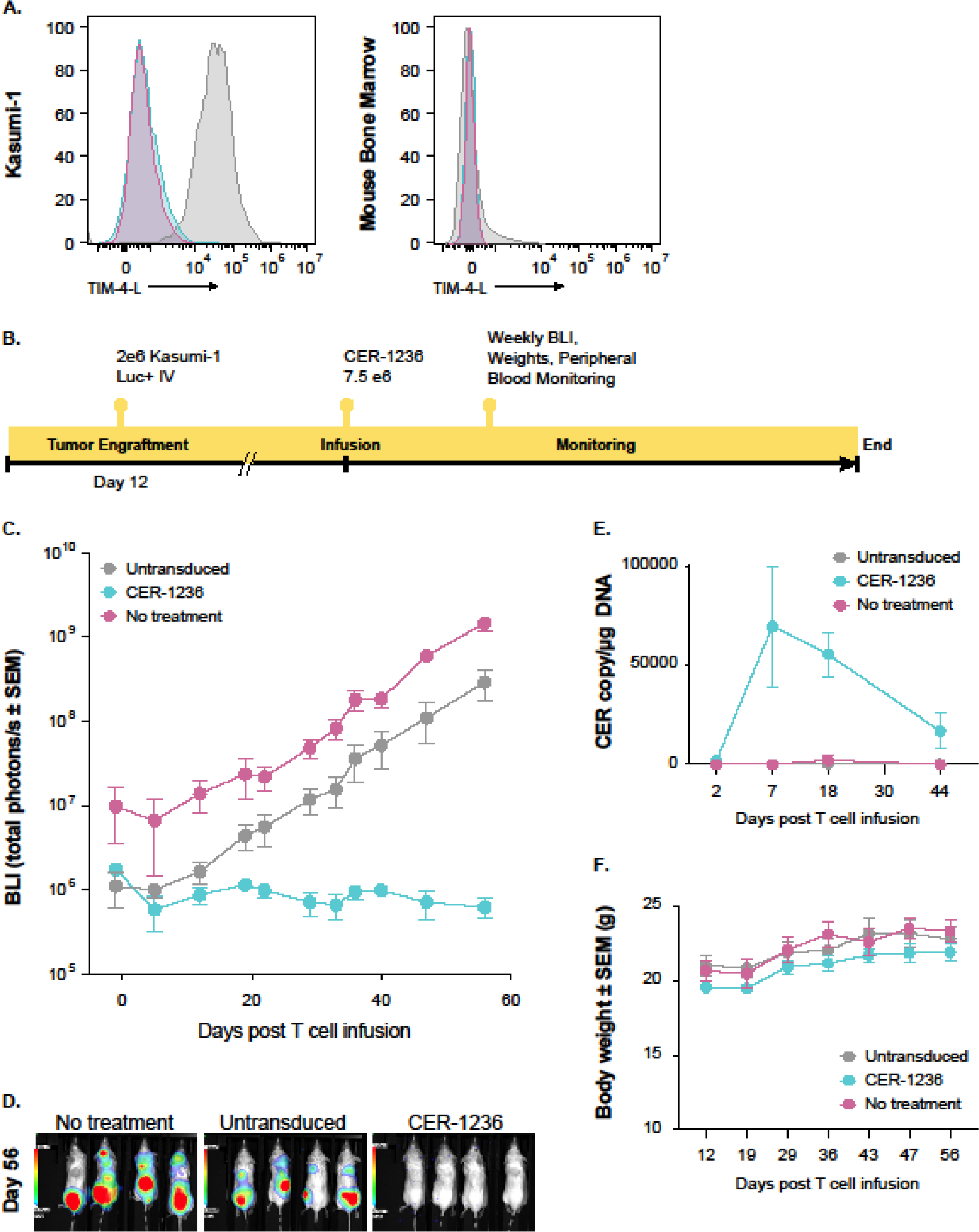
CER-1236 eliminates TIM-4-L-positive Kasumi-1 cells in vivo. (A) NSG MHC dKO mice were engrafted with 2×10^6^ Kasumi-1 Fluc^+^GFP^+^ cells i.v. After 68 days of engraftment, mice were euthanized and TIM-4-L exposure on GFP^+^ cells was assessed by flow cytometry. PS exposure on healthy mCD45-GFP-cells was assessed as well. Representative histogram plots are shown of unstained (blue), fluorescence minus one (FMO, pink), and stained (grey) samples. N=3. (B) Schematic of in vivo experiments. NSG dKO mice were engrafted with 2×10^6^ Kasumi-1 Fluc^+^GFP^+^ cells i.v. on day -12. On day 0, 7.5×10^6^ CER^+^ CER-1236 T cells or an equivalent number of untransduced cells were infused i.v. into engrafted animals. Tumor BLI, mouse body weight, and T cell expansion in the peripheral blood were assessed weekly. (C) Anti-tumor effect of CER-1236. BLI was monitored 1-2x per week to assess tumor progression. Average BLI ± SEM are shown for each group, n=4 for no treatment, 4 for untransduced, 6 for CER-1236. (D) Representative BLI images of engrafted animals at d56. (E) CER-1236 expansion in peripheral blood. Blood was drawn from treated animals weekly and expansion of CER-1236 T cells was monitored by ddPCR. Average CER copies per μg of DNA is shown ±SEM, n=4 for no treatment, 4 for untransduced, 6 for CER-1236. (F) Body weight of treated animals. Body weight of all animals on study was measured weekly starting at 12 days post T cell inoculation. Average body weight ± SEM is shown. Statistical significance was determined using a two-way ANOVA with Tukey’s post-test. * = p < 0.05, **** = p < 0.0001, ns = not significant.

## Conclusions

Genetically modified T cells have emerged as a promising immunotherapy for treating B-cell leukemias; however, their effectiveness in AML is yet to be established. Given the limited number of AML-restricted antigens, current strategies aim to target broadly expressed antigens shared between normal and leukemic cells (i.e., CD33, CD123, FLT-3), which can lead to significant myelotoxicity (1, 3) and risk for infectious complications (1). Here we describe upregulation of TIM-4-L in primary AML samples across multiple molecular risk groups and recapitulate these findings in AML cell lines. We previously demonstrated upregulation of TIM-4-L in aggressive lymphoma primary samples as well as therapeutic targeting using TIM-4-L-directed T cell therapy in lymphoma models. Here, we report preclinical efficacy of TIM-4-L-directed T cells in eradicating AML cells across various levels of TIM-4-L density in vitro and in vivo without observed toxicity.

Unlike many other tumor associated antigens that are similarly expressed on cancer cells and their healthy counterparts, TIM-4-L is typically absent on the surface of healthy cells (26). However, many tumors upregulate TIM-4-L (10, 27), and this phenomenon can contribute to the establishment of a tumor-permissive microenvironment (8). TIM-4-L binds to a number of endogenous receptors that impede anti-tumor responses (28). Furthermore, increased TIM-4-L levels in advanced tumor microenvironments are linked with the induction of intratumoral DCs with immature phenotype with reduced antigen presentation capacity (29), suppression of T cell responses (30), increased IL-10, and fewer tumor infiltrating lymphocytes (31). These immunosuppressive functions are often accompanied by dysregulation of don’t-eat-me signals (32-35). The high relative expression of TIM-4-L on newly diagnosed AML samples suggests that these tumors may gain an advantage from the immunoregulatory effects of TIM-4-L. However, the exact role of TIM-4-L in the establishment and maintenance of AML warrants further examination.

Importantly, we show TIM-4-L directed T cells were effective at eliminating AML cell lines that represent AML of medium or low antigen density. Interestingly, when TIM-4-L upregulation was assessed on ex vivo Kasumi-1 samples, we observed an increase in TIM-4-L intensity as compared to cells cultured in vitro, suggesting that TIM-4-L levels on malignant cells may be sensitive to environmental cues. Across various high-risk molecular subgroups, TIM-4-L measurements were detected at notably higher levels compared to cell lines, including *TP53*-altered AML and other adverse risk molecular subgroups, which are among the myeloid malignancies with the poorest prognosis and highest unmet need. If antigen density proves to impact TIM-4 targeting efficacy, combination strategies with various agents within the AML treatment landscape have potential to augment TIM-4-L surface density. In agreement with this, the 4 patients in our dataset that were receiving various AML therapies had overall high frequency and were in the upper tertile for intensity of surface TIM-4-L. Furthermore, given that CER-1236 is resistant to azacytidine, this offers the possibility of combining with azacytidine both for salvage and in the frontline setting. This is particularly relevant for *TP53* mutated MDS/AML where azacytidine remains the standard of care (SOC), as addition of other SOC therapies have not been shown to improve outcomes (36, 37). Additional work is required to understand the effects of concurrent anti-leukemic therapy on TIM-4-L expression and to devise strategies to enhance T cell efficacy.

Finally, as one mechanism for leukemia escape from engineered T cell therapies involves antigen loss or downregulation, approaches that have the capacity to target multiple antigens and or adapt to rapid clonal evolution offer unique advantages. CER-1236 has previously been shown to elicit both direct cytotoxic and indirect-mediated killing by enhancing antigen presentation to tumor associated antigens (17). Importantly, in this study and previous models using CER-1236 T cells, we have observed potent in vivo anti-tumor activity without on-target off-tumor toxicity, despite the conservation of TIM-4-L in both mice and primates (17, 38). These multifaceted mechanisms for tumor control make chimeric TIM-4 targeting T cells a promising candidate for overcoming antigen-related challenges for rapidly evolving leukemias.

Taken together, the results obtained in these studies demonstrate TIM-4-L is a viable target for T cell therapy in AML. Further investigation into the role of TIM-4-L in AML pathogenesis and its potential as an anti-leukemic target for clinical development is warranted.

