## Supplemental Tables and Figures for "Therapeutic Targeting of TIM-4-L With Engineered T Cells for Acute Myeloid Leukemia"

**Supplementary Table 1. Individual patient characteristics**

**Supplementary Figure 1. CER-1236 T cell product characterization.** (A) Cell viability for CER-1236 product was measured during the manufacturing process using automated cell counter (NucleoCounter® NC-200™ ) on each day of manufacturing. (B) CER-1236 T cells show robust expansion by day 7 of the manufacturing process (C) Percent transduction based on the proportion of cells expressing TIM-4 was determined by flow cytometry on day 7 of production, percentage of TIM-4$^{+}$ cells was determined among live, single, CD3$^{+}$ cells. (D) Genomic DNA was isolated from bulk CER-1236 cells on day 7 and VCN was measured using probes against the transgene and the endogenous gene histone acetyltransferase. To determine copy number per transduced cell, VCN obtained from bulk cell DNA was normalized to the frequency of TIM-4+ cells using the following equation: (bulk VCN / %TIM-4$^{+}$). (E) PD-1 levels are not altered during cell production. Levels of PD-1 at the start of cell production (day 0) and the end of cell production (day 7) were measured by flow cytometry. (F) The T-cell memory phenotype profile for CER-1236 T cells were characterized using flowcytometry on day 0 and day 7 of the manufacturing process. T-cells populations with expression of leukocyte common antigen isoform (CD45RA) and chemokine receptor (CCR7) are categorized as follows: CCR7+CD45RA+ = naïve, CCR7+CD45RA– = central memory, CCR7–CD45RA– = effector memory, and CCR7–CD45RA+ = terminally differentiated effector memory (TEMRA). (G) CER-1236 products were measured for product purity during manufacturing process using 8-color immunophenotyping panel, CD3+ T cells were further gated based on the CD4$^{+}$ and CD8$^{+}$ expression and CD4:CD8 T-cell ratio was obtained by diving % CD4 cells by % CD8 T cells within live, single, CD3$^{+}$ cells. (H) IFN-γ secretion in response to immobilized PS. IFN-γ secretion was measured by automated ELISA using supernatant of cells harvested on day 7 of the CER-1236 manufacturing process. Data is representative of 4 donors. Abbreviations: CER = chimeric engulfment receptor; VCN = Viral copy number; PD1 = Programmed cell death protein

**Supplementary Figure 2. Flow gating strategy for TIM-4-L detection on primary bone marrow.** TIM-4-L was assessed on bone marrow aspirates using flow cytometry. Samples were gated for FSCxSSC, singlets were gated using FSC-HxFSC-W, live cells were selected using Live/Dead Aqua negative. Within live cells, healthy cells were selected using CD45hi and AML cells were selected using CD45int. Gates were determined for each sample using FMO stain controls.

**Supplementary Figure 3. TIM-4-L in AML risk groups.** AML samples were categorized by ELN 2022 risk stratification into adverse, intermediate, and favorable risk. (A) TIM-4-L positivity and (B) TIM-4-L gMFI were determined. For adverse, intermediate and unknown risk, average is shown, ± SEM. For favorable risk, average ± range of duplicate measurements is shown.

**Supplementary Figure 4. TIM-4-L in peripheral blood mononuclear cells (PBMC).** 10 primary AML PBMC and 8 healthy donor PBMCs were stained for TIM-4-L using recombinant human TIM-4. (A) Percent positive TIM-4-L cells. Percent TIM-4-L-positive cells were identified on live, CD45dim cells. Gating was determined using TIM-4 FMO controls for each sample. Black dots indicate samples receiving therapy. Each sample was stained in duplicate and average is shown. (B) gMFI of TIM-4-L staining. The gMFI of TIM-4-L staining was measured on live, CD45dim cells. Gating was determined using TIM-4 FMO controls for each sample. Black dots indicate samples receiving therapy. Statistical significance was tested using student's unpaired t-test. ** = $p < 0.01$, ns = not significant.

**Supplementary Figure 5. Treatment of AML with 5-Azacytidine induces TIM-4-L.** Kasumi-1 or MV-4-11 AML cells were treated with various concentrations of 5-azacytidine and TIM-4-L exposure was measured after culture. (A) TIM-4-L exposure on Kasumi-1 cells. TIM-4-L exposure was measured at 120h of culture in the presence or absence of the indicated concentrations of 5-Azacytidine. (B) TIM-4-L exposure on MV-4-11-1 cells. TIM-4-L exposure was measured at 48h of culture in the presence or absence of the indicated concentrations of 5-azacytidine. Average of duplicate measurements is shown, ± SEM.

Supplementary Table 1. Individual patient characteristics

| Treatment Status | Prior Treatments | Patient Age At Collection | Gender | % Blast Cells | Risk Category |
|---|---|---|---|---|---|
| Newly Diagnosed | | 67 | Female | 91 | Adverse |
| Newly Diagnosed | | 59 | Female | 35 | Adverse |
| Newly Diagnosed | | 69 | Male | 75 | Intermediate |
| Newly Diagnosed | | 59 | Male | 93 | N/A |
| Newly Diagnosed | | 62 | Female | 30 | Adverse |
| Newly Diagnosed | | 71 | Male | 95 | N/A |
| Newly Diagnosed | | 48 | Male | 82 | Adverse |
| Newly Diagnosed | | 51 | Male | 80 | Adverse |
| Newly Diagnosed | | 38 | Female | 90 | N/A |
| Newly Diagnosed | | 74 | Male | 81 | N/A |
| Newly Diagnosed | | 62 | Female | 82 | Intermediate |
| Newly Diagnosed | | 43 | Male | 91 | Intermediate |
| Newly Diagnosed | | 85 | Female | 70 | N/A |
| Newly Diagnosed | | 69 | Female | 84 | Intermediate |
| Stable | Azacytidine | 71 | Female | 73 | Intermediate |
| Newly Diagnosed | | 41 | Male | 77 | Favorable |
| Progressive | Imatinib | 63 | Female | 50 | Intermediate |
| Newly Diagnosed | | 63 | Female | 83 | Adverse |
| Newly Diagnosed | | 83 | Male | 57 | Intermediate |
| Newly Diagnosed | | 72 | Female | 76 | Adverse |
| Newly Diagnosed | | 67 | Female | 15 | Intermediate |
| Newly Diagnosed | | 31 | Male | 86 | Intermediate |
| Newly Diagnosed | | 21 | Female | 95 | Adverse |
| Progressive | 7+3, Cytarabine | 51 | Male | 93 | Adverse |
| Newly Diagnosed | | 71 | Female | 87 | Adverse |
| Newly Diagnosed | | 33 | Female | 73 | Adverse |
| Newly Diagnosed | | 88 | Female | 92 | Adverse |
| Newly Diagnosed | | 46 | Male | 94 | Adverse |
| Newly Diagnosed | | 42 | Female | 76 | Adverse |
| Newly Diagnosed | | 66 | Male | 61 | Adverse |
| Newly Diagnosed | | 38 | Male | 74 | Adverse |
| Newly Diagnosed | | 44 | Female | 74 | Adverse |
| Progressive | Azacytidine | 85 | Male | 87 | Adverse |

Supplementary Figure 1

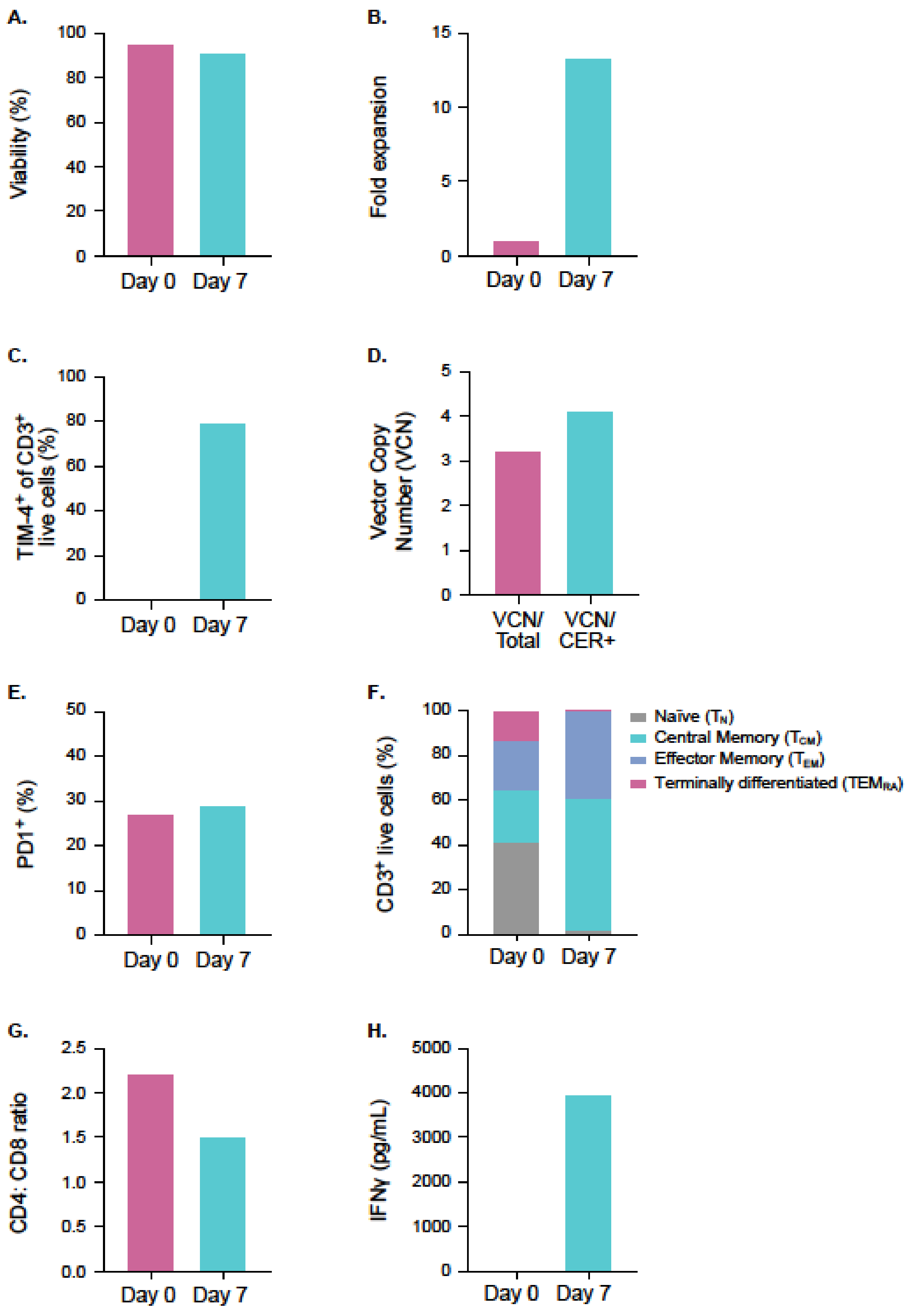

Supplementary Figure 2

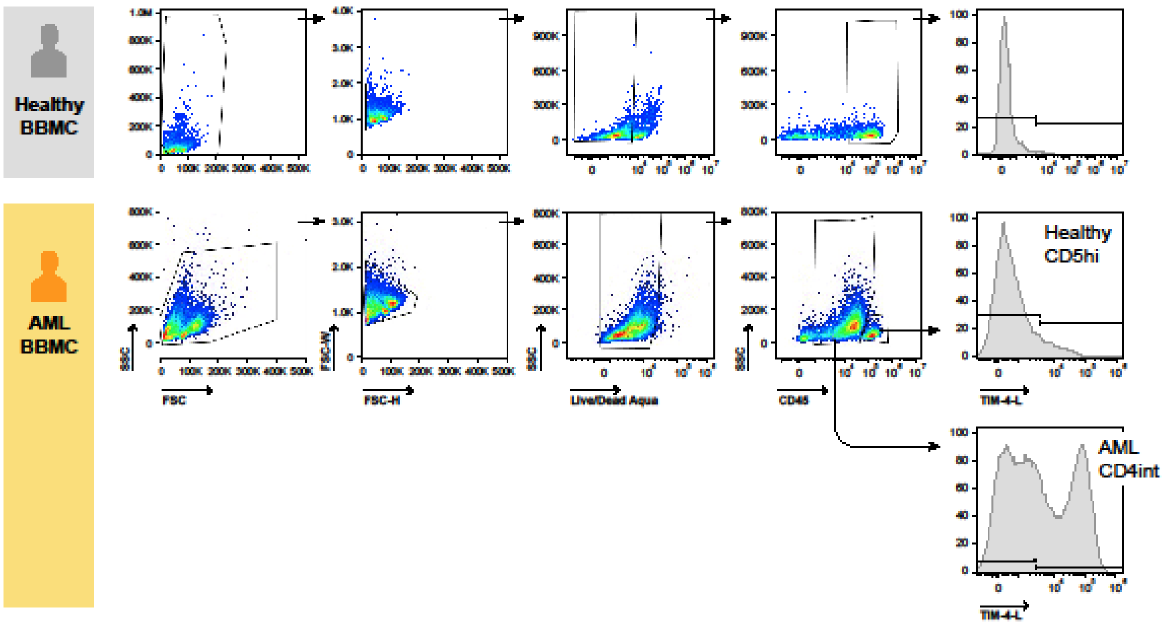

Supplementary Figure 3

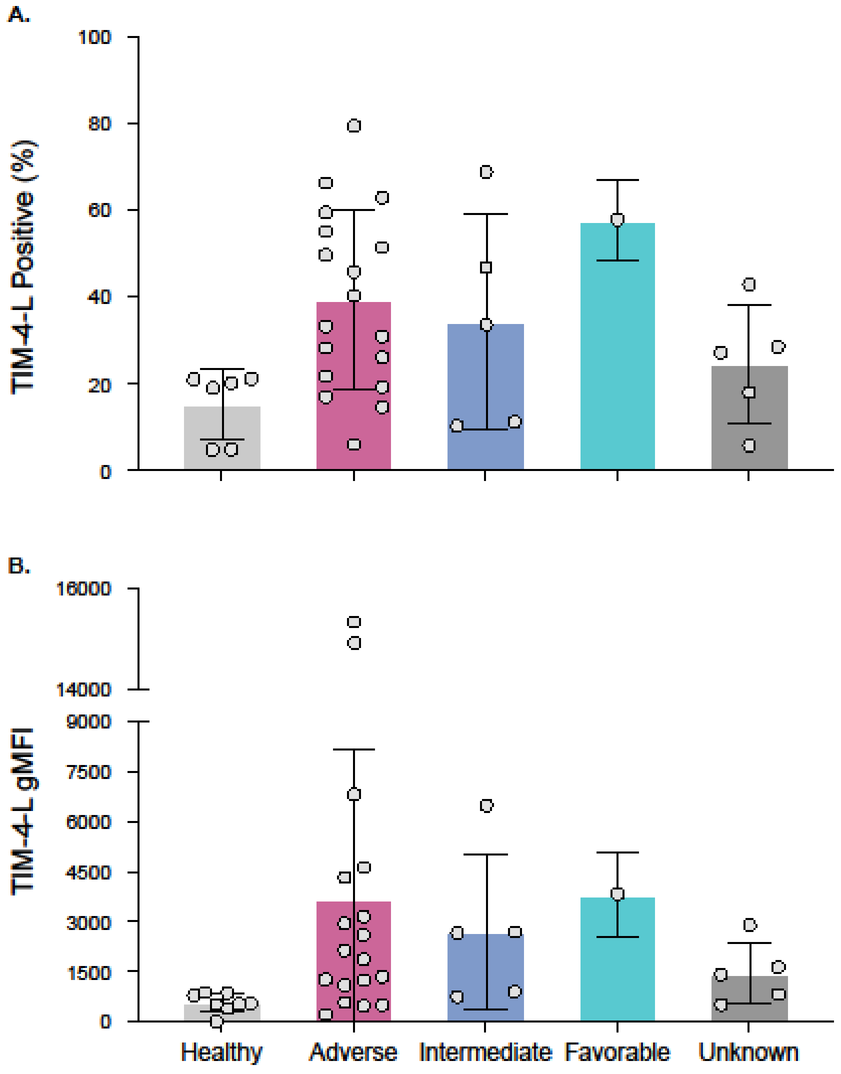

Supplementary Figure 4

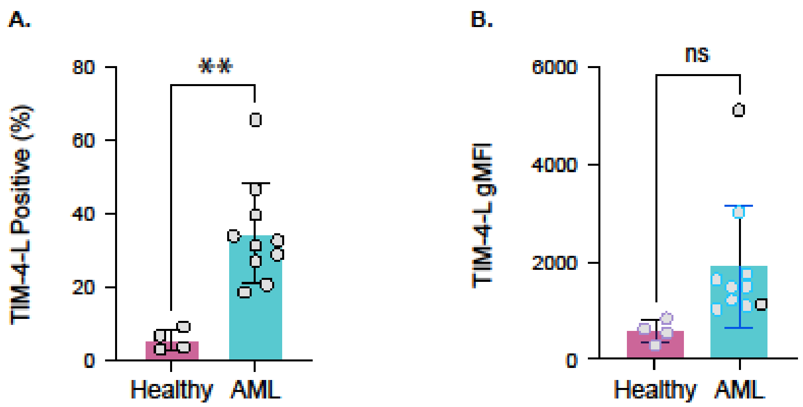

Supplementary Figure 5

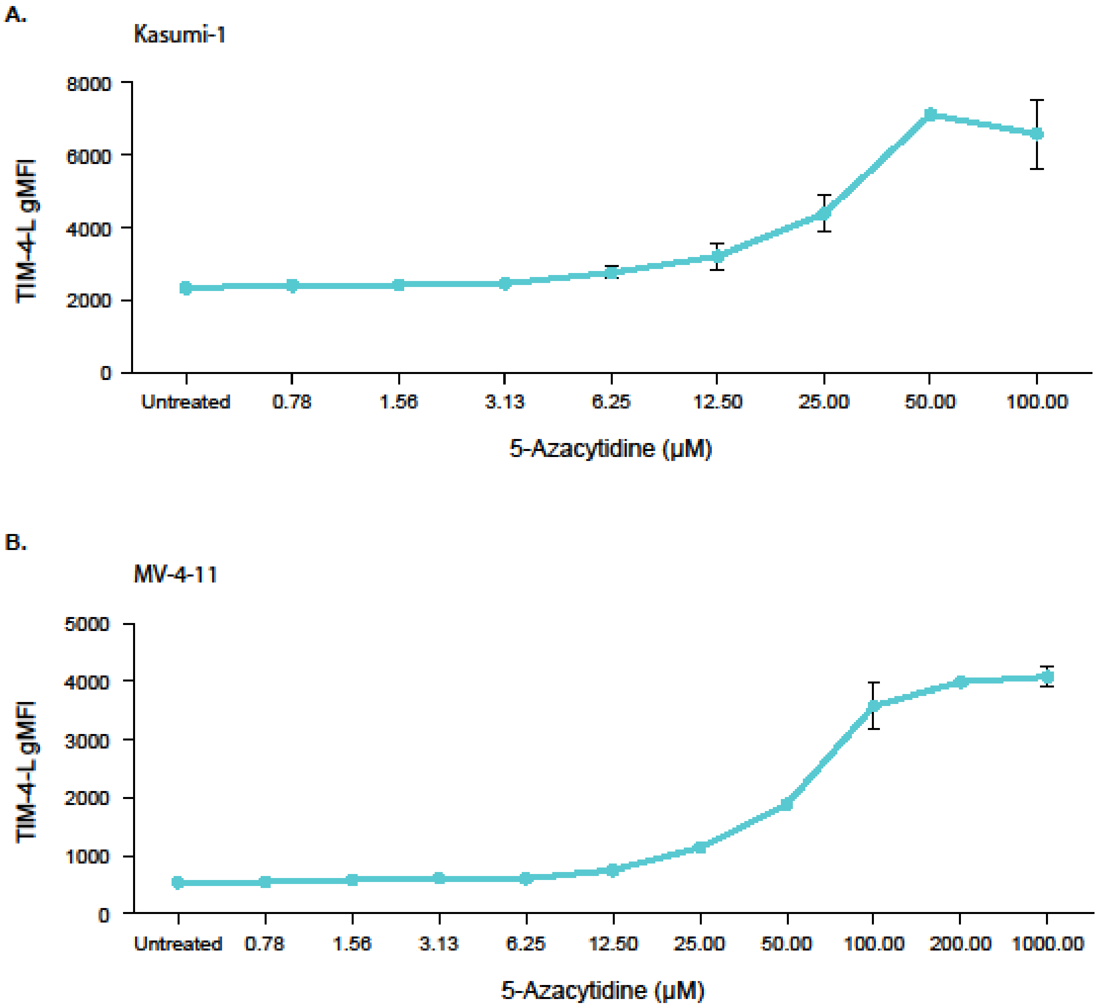
